# Evolution of mouse circadian enhancers from transposable elements

**DOI:** 10.1101/2020.11.09.375469

**Authors:** Julius Judd, Hayley Sanderson, Cédric Feschotte

**Affiliations:** Department of Molecular Biology and Genetics, Cornell University, Ithaca, New York 14853, USA; Department of Human Genetics, University of Utah School of Medicine, Salt Lake City, Utah, 84112, USA

**Keywords:** Transposable elements, enhancers, transcription, gene regulation, regulatory evolution, circadian rhythms

## Abstract

**Background:** Transposable elements are increasingly recognized as a source of cis-regulatory variation. Previous studies have revealed that transposons are often bound by transcription factors and some have been co-opted into functional enhancers regulating host gene expression. However, the process by which transposons mature into complex regulatory elements, like enhancers, remains poorly understood. To investigate this process, we examined the contribution of transposons to the cis-regulatory network controlling circadian gene expression in the mouse liver, a well-characterized network serving an important physiological function.

**Results:** ChIP-seq analyses revealed that transposons and other repeats contribute ~14% of the binding sites for core circadian regulators (CRs) including BMAL1, CLOCK, PER1/2, and CRY1/2, in the mouse liver. RSINE1, an abundant murine-specific SINE, was the only transposon family enriched for CR binding sites across all datasets. Sequence analyses and reporter assays revealed that the circadian regulatory activity of RSINE1 stems from the presence of imperfect CR binding motifs in the ancestral RSINE1 sequence. These motifs matured into canonical motifs through point mutations after transposition. Furthermore, maturation occurred preferentially within elements inserted in proximity of ancestral CR binding sites. RSINE1 also acquired motifs that recruit nuclear receptors known to cooperate with CR to regulate circadian gene expression specifically in the liver.

**Conclusions:** Our results suggest that the birth of enhancers from transposons is predicated both by the sequence of the transposon and by the cis-regulatory landscape surrounding their genomic integration site. This model illuminates how transposition fuels the emergence and turnover of enhancers during mammalian evolution.

## Background

Change in gene regulation is an important mechanism underlying the emergence of new biological traits [1–5]. There is a substantial body of empirical studies illustrating how the addition, modification, or disappearance of cis-regulatory elements, such as enhancers, has driven the emergence of profound phenotypic changes throughout evolution [6–8]. Thus, there has been an intensifying effort over the past decade to better understand mechanisms underlying the evolution of enhancers and other cis-regulatory elements [4,9–12].

In the broadest definition, enhancers are short (100 bp–1 kb) DNA sequences that modulate transcription of target genes regardless of genomic orientation or distance, and are often bound by transcription factors (TFs) [13,14]. Recent advances in functional genomics enabled nearly unbiased mapping of enhancers and their associated TF binding sites (TFBSs) on a genome-wide scale and facilitated systematic studies of enhancer evolution across and within species [11,15–17]. Seminal comparative studies in mammals revealed a low level of conservation in the genomic location of enhancers relative to genes and their promoters [18–27]. For instance, Villar and colleagues found that nearly half of 20,000–25,000 active liver enhancers mapped in each of 20 mammalian species are lineage- or even species-specific, while almost all promoters active in the liver are conserved across most or all the species examined [28]. However, recent analyses demonstrated that deeply conserved enhancers often coordinate robust and essential gene expression programs while less conserved enhancers contribute plasticity and redundancy to gene regulatory networks [29–31]. While these studies point to the rapid turnover of enhancers during mammalian evolution, the mechanisms underlying the birth and death of enhancers are only beginning to be understood [10,11,26,32–35].

Transposable elements (TEs) represent an important source of new cis-regulatory elements, including enhancers. TEs account for a substantial amount of nuclear DNA and genetic variation in virtually all metazoans [36]. For example, between one and two thirds of all mammalian genomes thus far examined are recognizable as being derived from TE sequences [36–39]. These elements inserted at various times during mammalian evolution, ranging from highly decayed copies integrated >100 million years ago to recently integrated copies that may be species-specific or still polymorphic in the population [37–42]. Several studies have systematically examined the contribution of TEs to TF binding and the birth of cis-regulatory elements, and some general principles have emerged [43–47]. First, TEs contribute a substantial but widely variable fraction (~2-40%) of the TFBSs mapped for a given TF throughout the genome [48–52]. Second, TFBSs and cis-regulatory elements derived from TEs tend to be evolutionarily recent, and are restricted to specific species or lineages [34,50,53]. For example, ∼20% of OCT4 and NANOG binding sites were derived from lineage-specific TEs in humans and mice [20]. This may be explained by the fact that the majority of TEs in any mammalian genome are themselves lineage-specific: for example, 85% of mouse TEs are not shared with the human [40] and 35% are not even shared with the rat [54]. Third, not all TEs contribute equally: for any given TF there is generally one or a few TE families that account for a disproportionate fraction of binding sites relative to their frequency in the genome [20,44,46,48,50,51].

Multiple studies have now confirmed that different TE classes and families contribute TFBSs for different TFs in different mammalian species, and that these TE-derived TFBSs occasionally undergo exaptation to give rise to new host regulatory elements (reviewed in [44,47]). However, the mechanisms by which complex enhancers emerge from TEs remain poorly understood. For instance, it is unclear why specific TE families or copies are bound by a particular TF while closely related elements in the same genome are not [45,51]. The path by which individual TE copies are co-opted for regulatory purposes has been scarcely characterized [55], and the relative contributions of combinatorial sequence motifs pre-existing within TE or in the vicinity of their insertion site, have not been examined in detail. To address these and other poorly understood aspects of TE co-option in regulatory evolution, we chose to examine their contribution to the cis-regulatory network underlying circadian gene expression. The machinery responsible for the transcriptional control of circadian gene expression is deeply conserved and has been extensively characterized in the mouse liver, which provides a solid experimental framework against which the impact of TEs can be queried. The circadian clock also presents the relatively unique advantage of providing a particularly robust system to examine the binding of TEs by regulatory proteins, as circadian rhythms are maintained by a series of interconnecting feedback loops of paralogous TFs [56–58].

The primary feedback loop consists of six circadian regulators (CRs), of which two are transcriptional activators (BMAL1 and CLOCK) and four are transcriptional repressors (PER1, PER2, CRY1 and CRY2). During the day, BMAL1 and CLOCK form a heterodimer, which binds to a tandem pair of E-box motifs in distal and promoter regions of clock-controlled genes [59]. Among the direct targets of the BMAL1:CLOCK complex are the repressors PER1/2 and CRY1/2. Following translation, PER and CRY enter the nucleus and inhibit BMAL1:CLOCK mediated transcription, thereby decreasing their own transcription and generating a feedback loop essential to the maintenance of the clock period [60,61]. This model has recently been revised to reflect reports that BMAL1 acts as a pioneer factor and promotes rhythmic nucleosome removal, and that transcription promoted by CLOCK:BMAL1 is not homogeneously oscillatory [62]. It is proposed that CLOCK:BMAL1 binding rhythmically maintains a chromatin landscape which facilitates binding and transcriptional activation by other ubiquitous or tissue-specific transcription factors, including members of the nuclear receptor (NR) family [63]. Interactions between CRs and liver-specific NRs are thought to underlie liver-specific circadian regulation of metabolic processes such as glucose, cholesterol, and lipid metabolism [64–66]. The vast amount of data and knowledge available for circadian regulation in the mouse liver provides a solid paradigm to dissect the mechanisms underlying the contribution of particular TEs to this cis-regulatory network.

## Results

### Repetitive elements contribute CR TFBSs with weaker enhancer chromatin signatures than non-repetitive CR TFBSs

To investigate the contribution of DNA repeats to cis-regulatory elements entwined in the mammalian circadian gene regulatory network, we turned to a seminal collection of ChIP-seq experiments that reported the genome-wide binding profiles of the six core CRs in the mouse liver [67]. To utilize the most recent mouse genome annotation, we aligned raw sequencing reads (Table 1) to the mm10 genome assembly and used uniquely mapped reads to call peaks. Of the peaks identified with this approach, ~60%-92% overlapped with peaks originally reported [67] (Fig. S1A). Cognate sets of CLOCK and BMAL1 peaks overlapped by ~85% (Fig. S1A). The fraction of peaks that did not overlap between the two sets (based on their distance from each other) were largely low confidence peaks (Fig. S1B). The fold change over background of the re-called peaks correlated with the summit height of the original peaks (Fig. S1C). Overall, the total number of peaks called for each CR was in good agreement with those reported in the published analysis [67] (Fig. S1D). Together these findings validate this set of CR binding sites in the mouse liver.

**Table 1:** Sources of data used in this study.

| Data | SRA | Citation |
| --- | --- | --- |
| CLOCK, BMAL1, CRY1, CRY2, PER1, PER2 ChIP-seq | SRP014752 | [67] |
| DNase-seq, H3K27Ac ChIP-seq, Pol II ChIP-seq | SRP045509 | [68] |
| GRO-seq, RORα ChIP-seq | SRP044381 | [69] |
| RXR, LXR, PPARα ChIP-seq | SRP010657 | [70] |
| REV-ERBα, REV-ERBβ ChIP-seq | SRP009472 | [71] |
| High confidence CLOCK:BMAL1 binding sites | N/A | [63] |
| Tissue Specific BMAL1 binding sites | SRP132869 | [72] |
| Hepa 1-6 RNA-seq | SRP059935 | [73] |

We then categorized CR binding sites as <u>R</u>epeat-<u>A</u>ssociated <u>B</u>inding Sites (RABS) [52] if the ChIP-seq peak coordinates showed positional overlap of >50% with a repetitive element annotated by RepeatMasker, which was accessed from the UCSC genome browser and filtered for low complexity and simple repeats [74–76]. This analysis revealed that between 8.3%–14.3% of each CR peak set mapped within TEs or other repetitive elements (Fig. 1A). To corroborate these results, we turned a published set of stringent CLOCK:BMAL1 binding sites that was derived from the same ChIP-seq data but required occupancy by both CLOCK and BMAL1 and was categorized as rhythmic, arrhythmic, or not expressed based on the circadian nascent transcriptional output of the nearest gene [63]. Rhythmic peaks were further classified as transcriptionally in or out of phase respective to CLOCK:BMAL1 binding. Interestingly, when we intersected these peaks with the coordinates of repetitive elements as described above, we observed slightly lower percentages (6.3%–9.7%) of RABS (Fig. 1B), which led us to speculate that TE-derived CR binding sites might be less likely to have biological relevance than non-repeat-derived CR binding sites.

**Figure 1:**
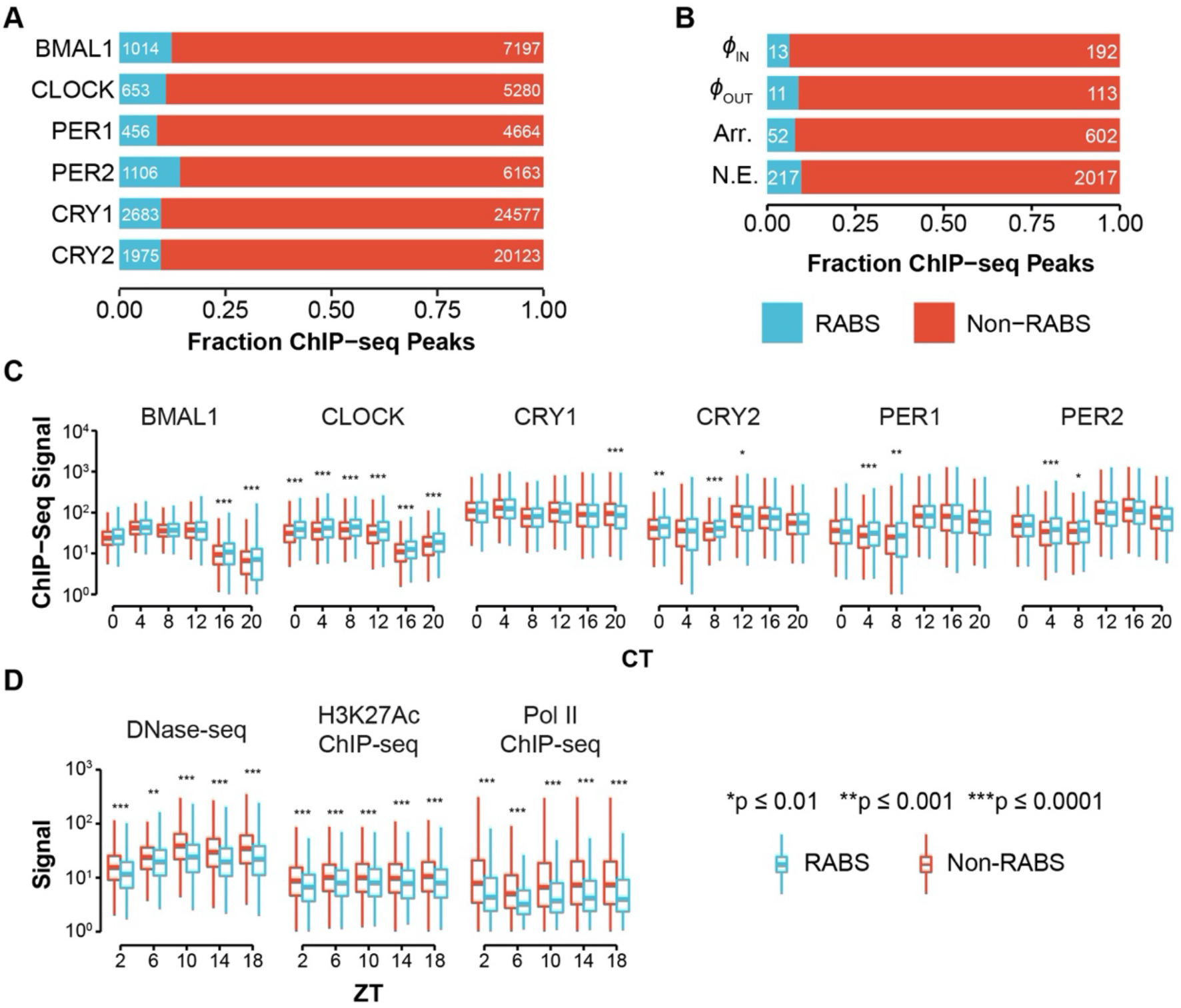
Repeat-derived CR binding sites display similar CR binding but less hallmarks of enhancer activity. **(A)** Fraction of ChIP-seq peaks for each CR for which >50% of the peak did (RABS) or did not (Non-RABS) map within repetitive elements. **(B)** CLOCK:BMAL1 ChIP-seq peaks sorted by the nascent transcriptional output of the closest gene (Rhythmic in-phase or out-of-phase; Arr: Arrhythmic; NE: Not Expressed), classified as RABS or Non-RABS as in **(A)**. **(C)** Distribution of ChIP-seq signal in RABS (n=1014) and Non-RABS (n=7197) BMAL1 ChIP-seq peaks across Circadian Time (CT). **(D)** Distribution DNase-seq, H3K27Ac ChIP-seq, and Pol II ChIP-seq signal in RABS (n=1014) and Non-RABS (n=7197) BMAL1 ChIP-seq peaks across Zeitgeber Time (ZT). Boxplots in **(C–D)** show the median (center line), the 1^st^ and 3^rd^ quartiles (hinges), and 1.5×IQR (whiskers). Significance values are from the Kruskal-Wallis test with Bonferroni-corrected Dunn post-hoc comparison.

To investigate the biochemical activity of RABS, we examined temporal genome-wide binding profiles of each CR. As a control, we selected a random set of repeats matching the familial composition of the RABS set. The binding pattern of CRs at RABS recapitulates that observed at Non-RABS: CLOCK and BMAL1 binding increases from CT0–CT8 and then tapers off, PER1/2 and CRY2 binding is initially low and increases from CT12–CT20, and CRY1 binding is maximally bound at CT0–CT4 but also increases at CT12 and CT20 (Fig.1C; Fig. S2A). When we quantified ChIP-seq signal within peaks, the only statistically significant difference consistent for all time points was that median CLOCK signal was increased in RABS relative to Non-RABS. However, the magnitude of this increase is subtle and led us to conclude that, overall, the oscillatory profile of CR binding at RABS and Non-RABS is essentially indistinguishable.

The preferred binding motif of the CLOCK:BMAL1 dimer is a pair of E-Box motifs (CACGTG) separated by a variable 6-7 nucleotide spacer [59]. We predicted that both RABS and Non-RABS would be enriched in E-Box motifs, and found that there was indeed a strong pattern of E-Box enrichment at the center of both RABS and Non-RABS peaks, but no such enrichment at the center of our set of randomly selected repeats (Fig. S3A). The average distance from RABS peaks and repeats to the closest E-Box was similar to that observed from Non-RABS peaks, and both sets are much closer than expected by chance (Fig. S3B). Of E-Box motifs associated with RABS, 82.5% fell within the boundaries of the associated repeat, indicating that the repeats themselves are contributing the binding site and are directly bound by CRs.

Because RABS are bound by CRs in a similar manner to Non-RABS, we would expect that they would have similar chromatin state and regulatory potential. To interrogate chromatin state and regulatory activity or CR binding sites, we used published H3K27Ac ChIP-seq, DNase-seq, and Pol II ChIP-seq experiments conducted in the mouse liver in Zeitgeber time [68], which is distinct from Circadian time in that animals are harvested over a light-dark cycle rather than constant dark conditions. We accessed raw sequencing reads (Table 1) and processed them with the software pipeline used for CR ChIP-seq data with minor adjustments (see methods). Intriguingly, while the oscillatory pattern of DNase sensitivity, H3K27 acetylation, and Pol II occupancy is maintained between RABS and Non-RABS, the median observed signal intensity is significantly lower at RABS than at Non-RABS (Fig. 1D). We confirmed this observation by visualizing individual loci as heatmaps (Fig. S2B-D). The magnitude of these differences between RABS and Non-RABS is substantially greater than those observed in CR binding (compare Fig. 1C and Fig. 1D). This could be a result of differences in the unique mappability of repetitive regions between experiments, but the DNase-seq, H3K27Ac ChIP-seq, and Pol II ChIP-seq libraries were sequenced with longer read length than the CR ChIP-seq libraries, so the unique mappability of these libraries should be better, not worse.

We then used the Genomic Regions Enrichment of Annotations Tools (GREAT) [77] to examine whether the genes proximal to E-Box motifs within RABS and Non-RABS ChIP-seq peaks (two nearest genes within 1 Mb) are enriched for particular mouse phenotypes. GREAT determines enrichment by comparing the gene associations of input sites with those of a user-defined background. When we used the entire mouse genome as background, RABS and Non-RABS displayed significant enrichment for similar annotations corresponding to fairly broad ontologies, which are largely related to liver function or circadian phase as previously reported [67] (Fig. S4A-B). We reasoned that these enrichments may be driven by the general open chromatin landscape of liver cells rather than by biological processes more specifically under circadian control in the liver. Indeed, when we repeated the GREAT analysis using all open chromatin regions—defined by DNase accessibility at any time point—as background, a different trend emerged. While Non-RABS were again enriched in phenotypes likely to be related to general liver function and circadian phase (Fig. S4C), RABS were enriched in categories related to pigmentation phenotypes, including abnormal digit pigmentation, non-pigmented tail tip, and variable body spotting (Fig. S4D). Collectively these analyses indicate that RABS and Non-RABS share similar CR occupancy and motif composition, but RABS exhibit weaker signals of regulatory activity as a whole and are tied to genes whose expression patterns could be related to species-specific phenotypic variation.

### RABS are enriched for RSINE1 elements

To determine if particular TEs were contributing CR RABS more often than expected from their frequency in the genome, we tested for enrichment of individual mouse TE families for their intersection with ChIP-seq peaks using a previously described pipeline [78]. We found that several TE families were significantly enriched in each CR dataset, but after imposing a cutoff of >50 observed bound copies per family, only a single family, RSINE1, stood out across 5/6 ChIP-seq datasets (Fig. 2A). While RSINE1 did not meet our threshold of 50 elements bound in the PER1 ChIP-seq data with 43 unique elements bound, enrichment of RSINE1 was still statistically significant. For each set of CR binding sites there were between 43 and 242 RSINE1-derived RABS, representing 3.2 to 5.4-fold enrichment over expectation (Fig. 2B). Importantly, other related B4 SINE families including B4, B4A, and ID_B1 were only significantly enriched for binding of 1-3 of the CRs and no consistent trend was apparent. In total, 328 unique RSINE1 elements in the genome were bound by at least one CR. Application of this method to CLOCK:BMAL1 peaks sorted by the nascent transcriptional output of the nearest gene [63] revealed similar levels of RSINE1 enrichment among ‘arrhythmic’ and ‘not expressed’ classes of binding sites, but no significant increase in RSINE1 elements in ‘rhythmic in-phase’ transcriptionally cycling binding sites (Fig. 2C). No RSINE1 elements were associated with ‘rhythmic out-of-phase’ transcriptionally associated binding sites. This corroborates our previous speculation that TE-derived CR binding sites are less likely to have biological relevance than non-RABS CR binding sites.

**Figure 2:**
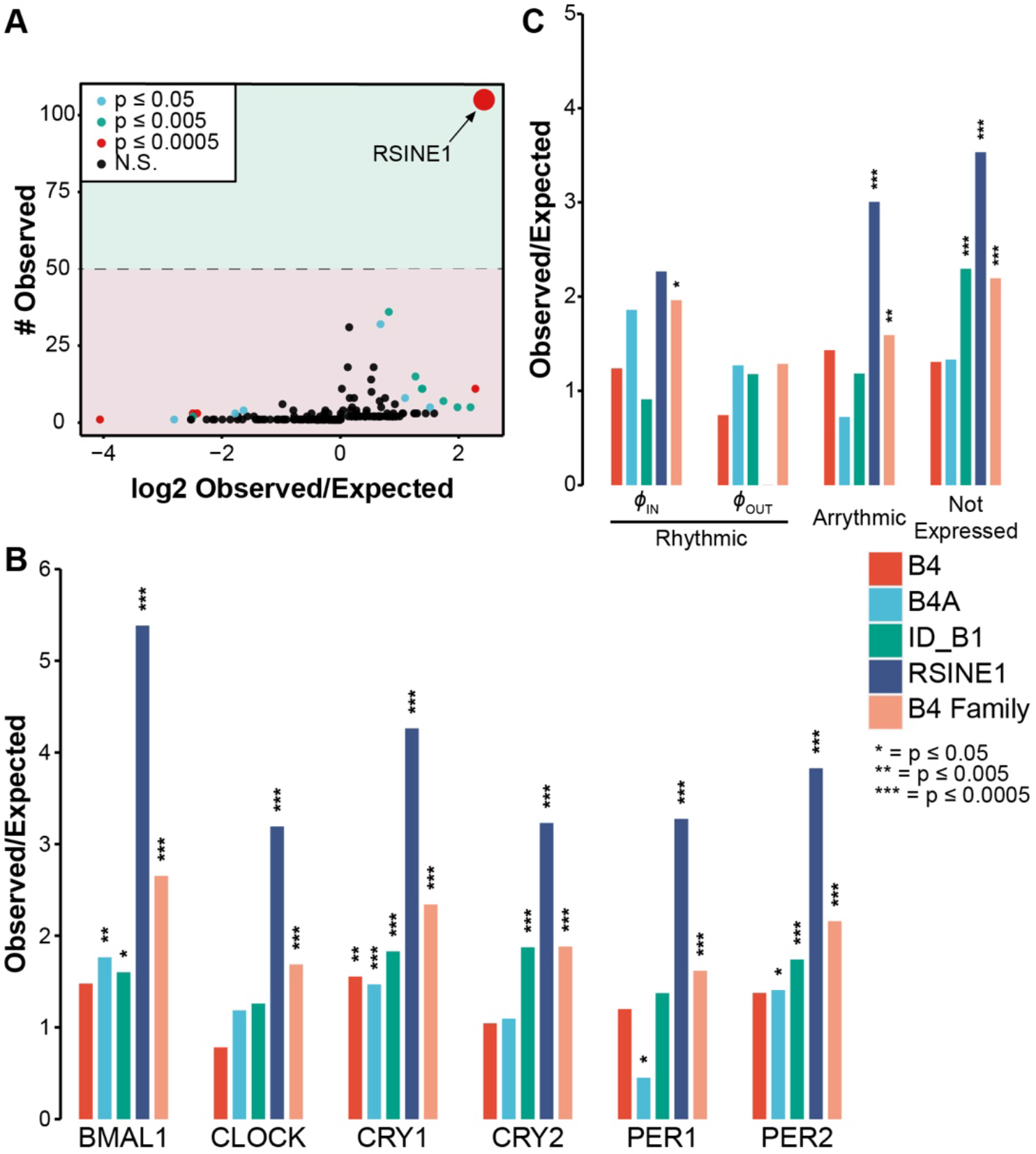
RSINE1 elements are overrepresented in RABS. **(A)** All TEs detected in BMAL1 ChIP-seq peaks, plotted by the number of elements observed compared to the ratio of observed to expected occurrences for that particular TE. Expected values were calculated by bootstrapping 1000 times (see methods). The dashed line denotes our cutoff of >50 elements observed imposed to filter false positives. **(B)** Ratio of observed to expected occurrences, as in **(A)**, for individual members of the B4 family and for the entire B4 family, in sets of RABS associated with each CR. **(C)** As in **(B)**, but enrichment of TEs in high confidence CLOCK:BMAL1 binding sites categorized by their nascent transcriptional output. All p-values were obtained using a two-sided binomial test.

To investigate why some RSINE1 elements are bound by CRs while others are not despite their identical sequence upon insertion, we first compared temporal CR binding patterns, chromatin landscape, and Pol II occupancy between RSINE1 elements predicted to be bound by CRs and those apparently unbound but still residing in open chromatin (OC; Unbound). We reasoned that RSINE1 elements in open chromatin regions might be more informative for comparison than random elements, though by doing so we are more likely to introduce false negatives in our unbound set. We found that there were strong oscillatory patterns of CR binding enriched at these bound RSINE1s, and while the pattern of enrichment on the unbound fraction was also oscillatory, it was weaker and more diffuse (Fig. 3A). We verified that this is only true of CR-unbound OC RSINE1s by inspecting heatmaps of unbound RSINE1s that do not reside in open chromatin and observed no detectable oscillatory CR ChIP-seq signal on these elements (Fig. S5). While this was somewhat surprising, we reasoned it could occur if these unbound OC RSINE1s were bound by other TFs themselves under circadian regulation or resided in genomic locale broadly regulated by CRs.

**Figure 3:**
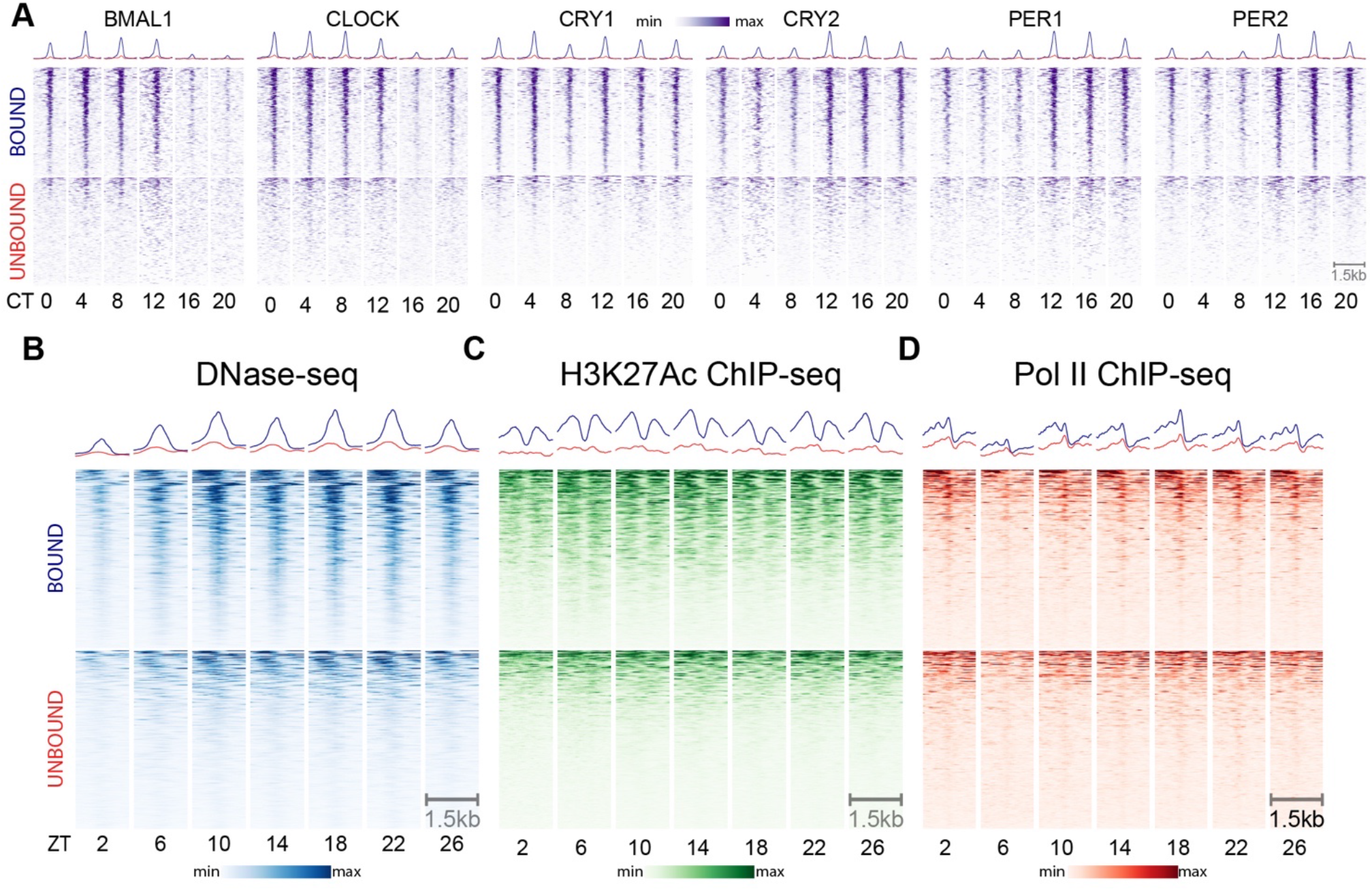
Differential CR occupancy, chromatin landscape, and Pol II occupancy of bound and unbound RSINE1 elements. **(A)** ChIP-seq signal for each CR across circadian time (CT), centered at CR associated RSINE1 elements (n=328, top) and a randomly selected set of 328 RSINE1 elements which fall within open chromatin regions as defined by DNase-seq but are not associated with CR binding (n=328, bottom). Heatmaps are centered at the middle of each element in positive orientation, extend 750 bp in each direction and are sorted by mean signal intensity across all rows for a given CR. **(B)** DNase-seq signal over Zeitgeber time (ZT). Regions, sorting, and centering as in **(A)**. **(C)** H3K27Ac ChIP-seq signal over Zeitgeber time (ZT). Regions, sorting, and centering as in **(A)**. **(D)** Pol II ChIP-seq signal over Zeitgeber time (ZT) Regions, s orting, and centering as in **(A)**. In all panels color scaling is relative to min-max signal within each block of heatmaps. Trace above heatmaps is the average signal in each class.

Furthermore, the vast majority of bound RSINE1s displayed patterns of DNase hypersensitivity (Fig. 3B), H3K27 acetylation (Fig. 3C), and Pol II occupancy (Fig. 3D) characteristic of active circadian enhancers. These patterns were rarely apparent, diffuse, and off-center relative to RSINE1 when examining unbound OC RSINE1s (Fig. 3B–D). While we expected that these OC RSINE1s would show DNase hypersensitivity—we selected this set precisely for this property—the oscillatory nature of the accessibility was surprising. We speculate this is driven by bound RSINE1s which did not meet the stringent 50% overlap criterion used in this analysis, and as such, our set of bound RSINE1s is likely an underestimate. Indeed, when we intersected our randomly selected set of unbound OC RSINE1 elements with BMAL1 ChIP-seq peaks, we found that 20/328 (6%) of these elements would have been classified as RABS if the overlap criterion was 25% instead of 50%. Taken together these results indicate that RSINE1 elements contribute a disproportionate fraction of CR TFBSs in the mouse liver, and that CR-bound RSINE1 elements display hallmarks of active regulatory elements.

### CR binding of RSINE1s is explained by sequence motif composition

We reasoned that the ancestral (consensus) sequence of RSINE1 might explain its propensity to bind CRs and promote the emergence of circadian enhancers. RSINE1 is a tRNA^Lys^-derived SINE family that belongs to the B4 superfamily, and its closest family relatives in the mouse genome are the B4 and B4A families [40,79,80]. While some other members of the B4 superfamily are significantly enriched to varying degrees in CR binding sites, none show levels of enrichment comparable to that of RSINE1 (Fig. 2B). Thus, we hypothesized that RSINE1 may have unique sequence motifs that predispose these elements to recruit CRs. To test this idea, we compared the consensus sequences of RSINE1, B4, and B4A obtained from Repbase [74] for the presence of E-box motifs (CACGTG), which are recognized by CLOCK and BMAL1, as well as RORE motifs (RGGTA), which recruit nuclear receptors (NRs). NRs are known to be important co-regulators of circadian gene expression in the liver and include RORα [81], REV-ERBs [82], and LXR/RXR [83], which have well characterized roles in liver metabolism [71] and circadian transcriptional regulation [57,66]. We also searched for D-box motifs, which are bound by TEF, HLF, and DBP; three additional circadian TFs that orchestrate rhythmic transcription in different phases [57].

We identified three E-Box motifs and two RORE motifs in the RSINE1 consensus sequence (Fig. 4A). All of the E-box motifs and two of the three RORE motifs present were not perfect matches to the empirically determined, optimal motifs for binding of CRs and NRs. Hereafter we call these imperfect motifs “proto-motifs”. A pair of closely spaced (6-7 nt) E-Box motifs (CACGTG) is the optimal CLOCK:BMAL1 heterodimer binding site [59]. We observed one E-Box that is a single nucleotide change (CAC<u>A</u>TG to CAC<u>G</u>TG; E-Box 1; Fig 4A) away from the optimal sequence in the center of the RSINE1 consensus sequence; however this motif is less likely to contribute significant regulatory activity as it is not a tandem motif, and similar sequences were present in B4 and B4A consensus sequences. Near the end of the RSINE1 consensus sequence, there were two E-Box motifs separated by 6 nt that are each a single nucleotide change away from the optimal sequence (CACG<u>C</u>G to CACG<u>T</u>G; E-Box 2 & 3; Fig. 4A). These two ancestral motifs each would require a single C to T mutation—a common deamination mutation in methylated DNA—to match the preferred CLOCK:BMAL1 binding site. The first RORE motif in the RSINE1 consensus is a perfect antisense motif and is closely followed by a sense motif that is two nucleotides away from an optimal RORE motif. However, these motifs are also present in the B4 and B4A consensus sequences. Together, these observations led us to conclude that E-Boxes 2 & 3 are the unique property of RSINE1 which most likely predisposed it to circadian regulatory activity.

**Figure 4:**
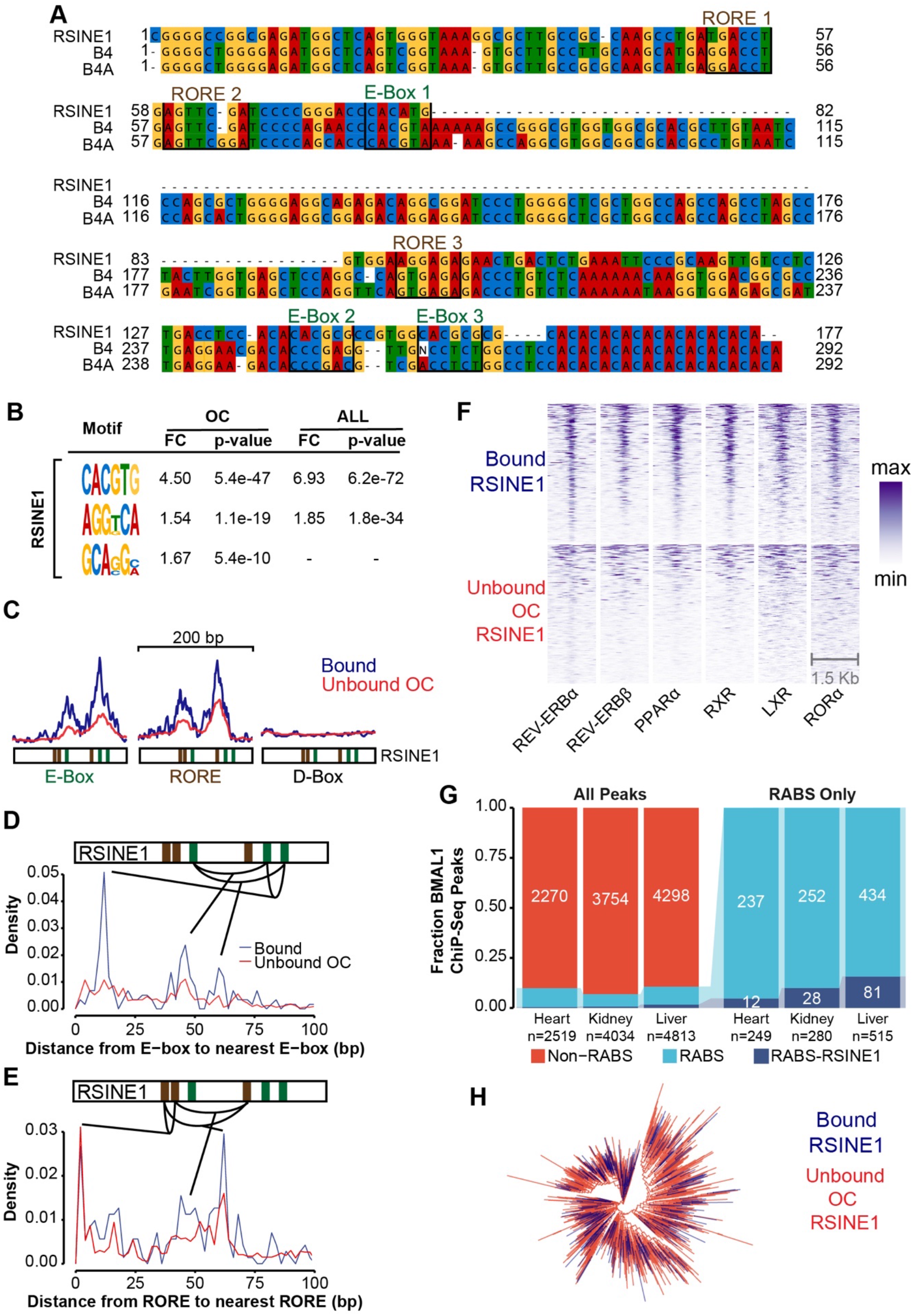
RSINE1 elements contain unique motif composition. **(A)** Multiple sequence alignment of B4 family SINES. RORE and E-Box pre-motifs are highlighted. **(B)** Motifs enriched in bound RSINE1 elements relative to unbound RSINE1 elements which reside in open chromatin (OC), or relative to all RSINE1 elements (ALL). The motif enrichment calculation was also performed using 500 bp of flanking sequences. **(C)** Density of D-Box (TTATGYAA), E-Box (CACRTG), and RORE (RGGTCA) motifs within bound (blue line; n=328) and unbound OC (red line; n=4279) classes of RSINE1 elements. Plots are centered on RSINE1s and extended 100 bp in either direction. **(D)** Average density of the distance from each E-Box and the next closest E-Box in bound RSINE1 elements (n=328) and unbound open chromatin RSINE1 elements (n=4279). **(E)** As in **(D)**, but density of the distance from each RORE to the next closest RORE motif. **(F)** ChIP-seq signal strength for various NRs centered at CR bound RSINE1 elements (n=328) or a randomly selected subset of unbound OC RSINE1 elements (n=328). Sort order is maintained across columns; color is scaled relative to min-to-max for each individual factor (column). **(G)** Fraction of BMAL1 ChIP-seq peaks in the mouse heart, kidney, and liver which are (RABS) or are not (Non-RABS) associated with repetitive elements by >50% overlap (left panel). Fraction of RABS which are or are not associated with RSINE1 elements (right panel) in each tissue. **(H)** Maximum likelihood tree of all CR bound RSINE1 elements (n=328) and a randomly selected set of unbound RSINE1 elements (n=1000) (see methods).

We next sought to understand why a subset of RSINE1 copies in the genome are bound by CRs while most are not. We searched for motifs enriched in CR-bound RSINE1 elements with respect to OC RSINE1s and all RSINE1s. Discriminative unbiased motif discovery returned CACGTG and AGGKCA as enriched in bound RSINE1 elements with respect to unbound RSINE1 elements. Strikingly, these motifs correspond to E-Box and RORE elements respectively (Fig. 4B). To further explore the motif composition of these elements, we plotted profiles of E-Box and RORE motif occurrence weighted by motif confidence over 200 bp regions around the center of bound and OC unbound RSINE1s (Fig. 4C). Bound and unbound fractions both displayed characteristic patterns of tandem E-Box and RORE motifs, but few D-Box motifs. Notably, the summary plot revealed that while the spatial pattern is similar, the bound fraction has higher signal strength, which indicated that bound RSINE1 elements more frequently contain motifs that more closely match the optimal sequence than unbound OC RSINE1 elements.

We next examined the density distribution of distance from each optimal E-Box motif and the next closest E-box motif in order to identify characteristic spacing, which might better explain the difference between bound and unbound RSINE1 elements. Strikingly, the bound class had a strong enrichment for an E-Box ~12 bp from the nearest optimal E-Box, which was not present in unbound OC RSINE1 elements (Fig. 4D) and coincides with the pair of proto-motifs at the 3’ end of the consensus (Fig. 4A). When we examined the distance from RORE to RORE, we observed enrichment of closely adjacent motifs in both classes of RSINE1 elements (Fig. 4E). These results are consistent with the spacing of proto-motifs in the RSINE1 consensus and support the model that the bound class of RSINE1s gained their regulatory activity after transposition via maturation of imperfect CR/NR binding sites.

To verify that identified RORE elements are actually bound by NRs in the mouse liver, we turned to published ChIP-seq data for REV-ERBs [71], RXR, LXR, PPARα [70], and RORα (Table 1). These datasets revealed that bound RSINE1 elements were occupied by the aforementioned NRs, and that different NRs were bound simultaneously at the same RSINE1 elements, indicating that different NRs and/or CRs could be binding to these elements cooperatively. A randomly selected subset of unbound RSINE1s in open chromatin also displayed some NR binding but not to the degree of CR bound RSINE1s (Fig. 4F). We reasoned that because NRs are liver-specific circadian co-regulators, the interplay of these CR/NR binding sites in the RSINE1 consensus may have given rise to tissue-specific regulatory elements. We then leveraged a recent analysis of tissue specific BMAL1 binding sites using ChIP-seq in the mouse liver, heart, and kidney [72], and found that while the fraction of peaks overlapping with repetitive elements was relatively consistent, the fraction overlapping with RSINE1 elements was highest in the liver and markedly lower in other tissues (Fig. 4G). These results suggest that the majority of RSINE1-derived CR binding sites are liver-specific.

To rule out the possibility that the E-Box and RORE motifs enriched in bound RSINE1s could have emerged from the expansion of a slightly different progenitor containing pre-existing optimal motifs but forming a distinct RSINE1 subfamily, we aligned all CR-bound RSINE1 elements located in open chromatin with 1000 randomly selected RSINE1 copies and performed a phylogenetic analysis, the resulting tree had star-like topology with no subfamily structure coinciding with the two categories (Fig. 4H). This confirms that both bound and unbound RSINE1 elements descend from the same ancestral sequence which amplified rapidly throughout the genome. These results led us to conclude that RSINE1 spread imperfect CR/NR binding sites throughout the mouse genome, providing fodder for the evolution of new circadian enhancers rather than spreading ‘ready-made’ enhancer modules.

### Circadian enhancer evolution from RSINE1 elements is context-dependent and lineage-specific

We next asked if the genomic location where RSINE1 inserted influences their propensity to act as circadian regulatory elements. More specifically, we asked whether the presence of an existing CR-bound sequence might favor the emergence of CR binding at RSINE1 elements. We determined the distance from each bound RSINE1 to the nearest non-repeat derived CR binding site and compared this to the expected distance using the same calculation but using all genomic copies of RSINE1. Bound RSINE1s are significantly closer to Non-RABS (median ≈ 8 kb) than expected given the genomic distribution of all RSINE1 elements (median ≈ 65 kb; Fig. 5A). To determine whether these Non-RABS existed prior to RSINE1 insertion and thus shaped the evolutionary trajectory of RSINE1, we then queried whether Non-RABS CR binding sites have deeper phylogenetic roots than those in associated with repetitive elements by determining the fraction of regions for which an orthologous region could be identified in the rat and human genomes using liftOver [84]. We found that ~46% of the Non-RABS ChIP-seq peaks could be traced to orthologous regions in the human genome, and ~74% to the rat genome (Fig. 5B). This percentage is higher than the overall percentage of the mouse genome (~40%) that can be aligned confidently at the nucleotide level with the human genome [40], indicating that Non-RABS tend to be deeply conserved. Consistent with RSINE1’s murine specificity, ~60% of bound RSINE1 elements could be traced to an orthologous region in the rat genome, but virtually none (0.3%) were detected in the human genome (Fig. 5C). Thus, in general, RSINE1 derived CR binding sites are evolutionarily younger than Non-RABS and must have inserted nearby existing CR binding sites prior to their maturation into circadian enhancers. Given our previous finding that RSINE1 elements contained proto-motifs prior to their expansion, this result suggests that RSINE1 elements preferentially matured into optimal circadian regulatory elements after insertion nearby existing circadian regulatory sequences.

**Figure 5:**
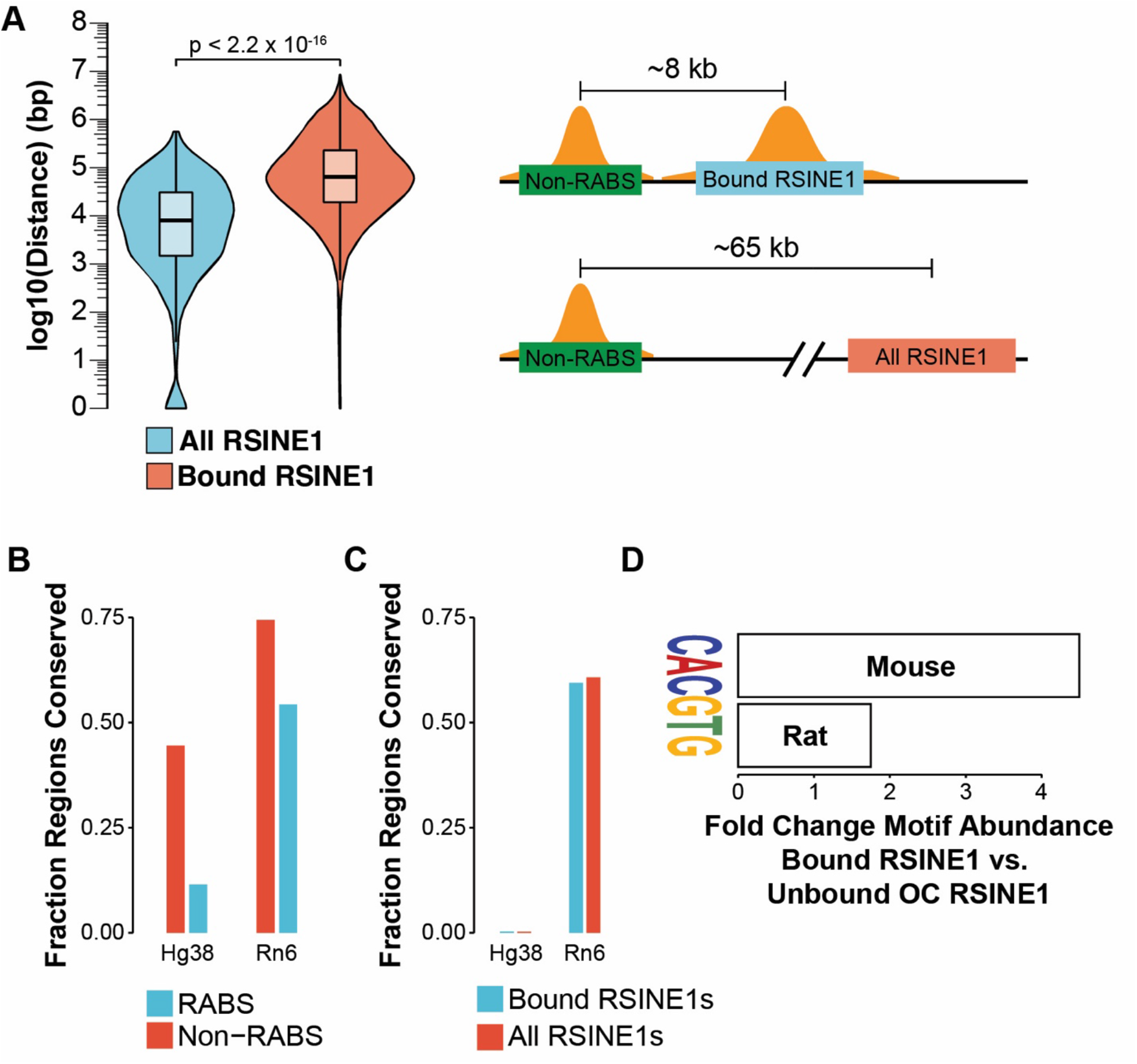
Evolution of CR binding sites from RSINE1 is context dependent. **(A)** Distribution of distance from each CR bound (n=328) or all (n=115,806) RSINE1 elements to the nearest non-repetitive CR binding site (Non-RABS). Boxplots show the median (center line), the 1^st^ and 3^rd^ quartiles (hinges), and 1.5×IQR (whiskers). Statistical testing was done using the Mann-Whitney/Wilcoxon rank sum test. **(B)** Fraction of Non-RABS and RABS CR bindings sites that can be traced to the human or rat genomes using liftOver. **(C)** Fraction of CR bound and unbound RSINE1 elements that can be traced to the human or rat genomes using liftOver. **(D)** Fold change CACGTG motifs detected in bound RSINE1 elements compared to unbound OC RSINE1 elements in the mouse as well as in syntenic orthologs in the rat.

RSINE1 elements appear to have distributed imperfect CR/NR binding sites prior to the divergence of the mouse and rat lineages, and we hypothesized that they occasionally matured into lineage- or tissue-specific circadian enhancers. To test this, we queried syntenic orthologs of mouse bound and unbound OC RSINE1 elements in the rat and performed motif discovery. This revealed that most elements which have developed perfect E-Box motifs (CACGTG) in the mouse have not developed the same motif in the rat (Fig. 6C). This indicates that bound RSINE1 elements in the mouse represent a subclass of RSINE1 elements which gained CR binding after the divergence of mouse and rat and thus are lineage specific CR binding sites. Together these findings show that the evolution of circadian regulatory elements from the proto-motif content of RSINE1 is a process which is context-dependent and lineage-specific.

**Figure 6:**
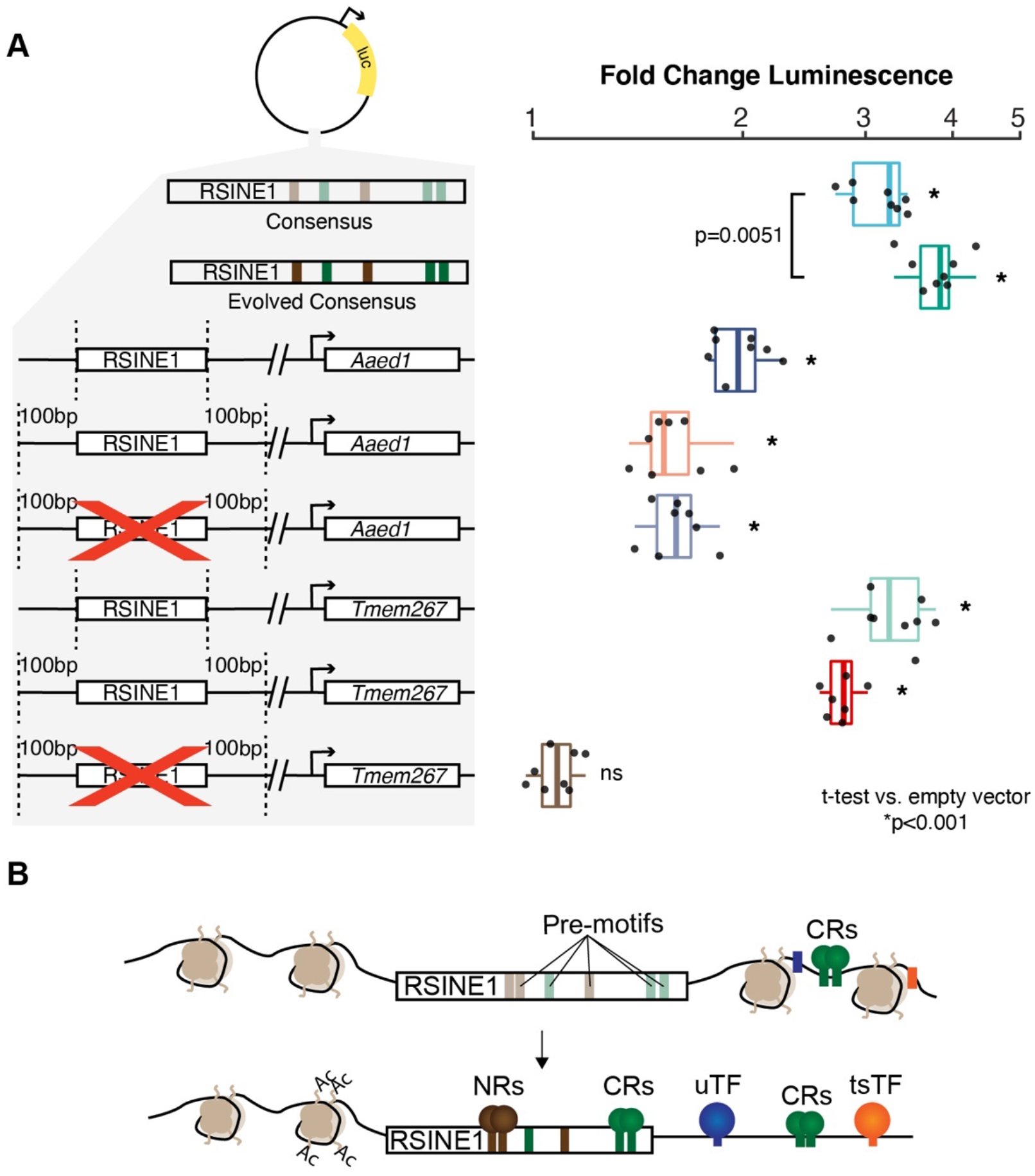
RSINE1 elements have enhancer activity *in vitro*. **(A)** Fold change luminescence when compared to an empty vector control after normalization to *Renilla* luciferase signal of enhancer reporter plasmids with various RSINE1-derived sequences inserted downstream of the luciferase gene. Included are: RSINE1 consensus as is or with “evolved” E-Box and RORE motifs at highlighted locations (all motifs were changed to the canonical motif with no other changes), an RSINE1 element upstream of the promoter of *Aaed1* (See Fig. S6B), and an RSINE1 upstream of the promoter of *4833420G17Rik* and *Tmem267* (See Fig. S6A). *Tmem267* and *Aaed1* RSINE1s were cloned as the RSINE1 alone, with 100 bp of flanking sequence, and with the RSINE1 deleted from the flanking region as a reconstruction of the ancestral state. Experiments are representative of two biological transfection replicates with four technical replicates each. Significance was assessed using pairwise two-tailed t-tests compared to the empty vector. **(B)** A model of how RSINE1 has fine-tuned the circadian regulatory landscape of the murine lineage by distributing imperfect CR/NR binding sites with moderate regulatory activities that modulate the activity of existing nearby circadian regulatory elements.

### A minority of RSINE1 elements have matured into circadian enhancers

The above data indicate that RSINE1 elements are poised for maturation into circadian regulatory elements. We next searched for evidence for circadian enhancer activity of RSINE1 elements. A published study [69] used GRO-seq (Table 1) to measure nascent transcription over circadian time in the mouse liver and identified enhancer RNAs (eRNAs) that transcriptionally oscillate. We intersected our set of CR-bound RSINE1 elements with the coordinates of these oscillating eRNAs and identified 37 RSINE1 elements (11.3% of bound RSINE1s) for which 50% of the RSINE1 element falls within the coordinates of an oscillatory eRNA. One such example is an RSINE1 element that falls within the last intron of *Paip1* and is ~6kb upstream of the start of *4833420G17Rik*, a cDNA with no identified function, which is in turn ~2kb upstream of the TSS of *Tmem267*, a gene with a CR-bound promoter (Fig. S6A). Furthermore, an independent study [63] identified *4833420G17Rik* as transcriptionally oscillating using Nascent-seq, and the RSINE1 element also overlapped with a high confidence BMAL1:CLOCK binding site designated as associated with rhythmic in phase transcription. Upon further inspection, we observed that the RSINE1 element had a characteristic pattern of oscillating CR binding and chromatin accessibility and was bound by NRs (Fig. S6A). When we examined the sequence composition of this RSINE1, we observed two E-Box motifs at the end of the element and a RORE motif in the center. A D-Box motif was also present slightly upstream of the element, indicating that this RSINE1 could have matured into a bona fide circadian enhancer by virtue of the genomic context it inserted into. Another RSINE1 that falls within an oscillatory eRNA but does not have a D-Box in the flanking sequence lies immediately upstream of the promoter of *Aaed1* (Fig. S6B).

We next used *in vitro* luciferase assays in Hepa 1-6 cells, a mouse hepatoma-derived cell line, to measure the enhancer activity of selected RSINE1 sequences. Importantly we verified that CLOCK and BMAL1 are constitutively expressed in this cell type using public RNA-seq data (Table 1). We initially compared the consensus sequence from Repbase [74] to an “*in silico* evolved” sequence that we subjected to a total of 7 nucleotide substitutions to perfect the degenerate motifs in the RSINE1 consensus. We observed that the consensus alone had moderate enhancer activity, and the “evolved” version’s activity was significantly boosted (Fig. 7A). Next, we examined the genomic RSINE1 elements described above (*Tmem267* and *Aaed1*). For each, we tested the regulatory activity of the RSINE1 element alone, the RSINE1 with 100 bp flanking sequences, and a proxy for the ancestral locus where the RSINE1 was removed leaving only the flanking sequences joined at the insertion site. In both cases, we found that the RSINE1 element alone had the highest activity, and that this activity was slightly lower when the flanking sequences were included (Fig. 6A). Removing the RSINE1 had no effect on the activity of the flanking sequences in the Aaed1 reporter, but completely abolished luciferase signal of the *Tmem267* reporter. This pattern was consistent with the presence of a D-Box motif in the upstream flank of the *Tmem267*, which is known to be bound by the transcriptional repressor E4BP3 (also called NFIL3) as part of a secondary circadian transcriptional feedback loop with opposing phase to the classic CLOCK:BMAL feedback loop [85]. The most likely explanation for the phenomenon observed at the *Aaed1* RSINE1 is that the flanking sequence contains both activating and repressive TFBSs which in balance yields mild basal enhancer activity, and while the RSINE1 contains stronger regulatory activity on its own, its presence in the flanking sequence does not disrupt this balance in a manner sufficient to alter enhancer activity. *These* assays illustrate how RSINE1 has shaped the circadian regulatory landscape of the murine lineage (Fig. 8) and highlight how the genomic context of each insertion of an identical family of repetitive elements dictates which elements gain TF binding and regulatory activity each in unique ways.

## Discussion

In this study, we show that a subset of CR binding sites in the mouse liver map within repetitive elements. This class had patterns of oscillatory CR binding that were indistinguishable from their non-repeat derived binding sites, but characteristics consistent with their function as bona fide circadian enhancers were weaker than those observed for non-repeat derived binding sites. This is in line with recent observations that only a small subset of TEs which display enhancer marks in mouse embryonic stem cells trigger gene expression changes when perturbed [86]. Furthermore, GO analysis of genes associated with TE-derived CR binding events show enrichment for genes linked to morphological phenotypes, including non-pigmented tail tip and variable body spotting, which are likely to be intra-specifically variable. We speculate that these associations reflect cis-regulatory connections that are evolutionarily young and might drive phenotypic plasticity akin to what is seen at the *Agouti* gene in mice, where the differential regulatory activity of IAP individuals lead to coat color variation [87–89]. Further study is needed to evaluate whether CR-bound TEs have directly contributed to changes in circadian gene expression with possible phenotypic consequences.

### Seeding and maturation of proto-motifs by transposition

In this study, we examined the properties that predispose one family of TEs, RSINE1, to be bound by CRs and to acquire the biochemical hallmarks of circadian enhancers. We focused on the RSINE1 family because it was the only TE family in the mouse genome that was significantly overrepresented for CR-binding events across all ChIP-seq datasets examined, suggesting that this family was in some way predisposed to be bound by CR. Furthermore, we observed that many of the CR-bound RSINE1 were also bound by NRs which are important regulators of circadian gene expression specifically in the liver [90,91]. In addition to being bound by CRs and NRs, a small subset of these RSINE1 elements displayed features characteristic of circadian enhancers, including oscillatory H3K27 acetylation, DNase accessibility, and nascent transcription of eRNAs. We found that the recurrent binding of CRs and NRs to RSINE1 elements can be explained in part by the presence of sequence motifs in their ancestral sequence that are close, but imperfect match to the E-Box and RORE motifs that recruit CR and NR, respectively. In particular, a feature unique to RSINE1 when compared to other closely related SINE families, is a pair of imperfect E-Box motifs at the 3’ end of the RSINE1 consensus sequence. Thus, unlike other cases of TE families enriched for TF binding events [20,45,48,50,51,92,93], RSINE1 did not introduce perfect TFBSs upon integration, but proto-motifs that repeatedly matured into optimal TFBSs within individual TE copies after their genomic integration.

Our analysis indicates that recurrent C-to-T mutations within the two proto E-boxes located at the 3’ end of RSINE1 represent the primary mutational path by which these elements acquired canonical CR binding sites. Interestingly, the C-to-T transition required within each of the proto-motif is at a CpG site (CACG<u>CG</u>-> CACG<u>TG</u>), which in mammalian genomes is ~10 times more likely to occur compared to mutations at non-CpG sites because of spontaneous deamination of methylated cytosine [94–96]. Because SINEs are heavily methylated in the mouse germline [97–99], it is likely that deamination at methylated CpG sites accelerated the maturation of the proto E-boxes residing within RSINE1 elements. The birth of TFBSs via CpG deamination in TEs has been observed previously in human Alu SINEs, which harbor proto-motifs for p53, PAX-6, and Myc that frequently mutate to introduce new binding sites for these factors [100,101]. Interestingly, Myc also binds an E-Box motif, though the proto-motif in the Alu consensus is CGCGCG, and thus requires two C-to-T transitions to mature into the canonical CACGTG motif. Our findings bolster the proposal that CpG deamination-mediated evolution of TFBSs from TE substrates is a pervasive force driving the emergence of new regulatory elements [100,101]. This may be especially common for motifs such as the E-box for which many possible configurations of proto-motifs can generate a canonical motif (CACGTG) via CpG deamination. This is notable because the E-box is the binding motif of multiple TFs with diverse physiological or developmental functions, including CLOCK:BMAL1, MYC, and MYOD.

### Genomic environment of TE insertions influences their regulatory trajectory

Our study of RSINE1 also underscores the role of the cis-regulatory context surrounding a TE insertion site in modulating the cis-regulatory potential of that TE. Our analyses indicate that RSINE1 elements that inserted nearby existing CR binding sites were predisposed to acquire CR-binding activity upon maturation of their E-Box proto-motifs. This observation suggests a scenario whereby the insertion of RSINE1 near pre-existing CR binding sites determined, at least in part, whether they would become bound by CRs. In other words, the genomic regions where CRs were already binding at the time of RSINE1 amplification represented a favorable environment for the recruitment of CR to RSINE1 inserted in these regions. It is also possible that RSINE1s were only permitted to develop *bona fide* CR/NR binding sites when their insertion location allowed this gain of activity to fine-tune, replace, or become redundant with existing circadian regulatory elements, while the introduction of such regulatory activity at other genomic locations might have been deleterious. Our reporter assays are consistent with the notion that the genomic sequences flanking CR bound RSINE1 elements temper their cis-regulatory activity. This is in line with the idea that RSINE1 elements that were permitted to mature into CR enhancers did so preferentially when inserted in genomic regions that buffered the deleterious potential of aberrant circadian regulatory activity.

Our observation that RSINE1 elements preferentially gained CR binding activity when located nearby existing CR BS is reminiscent of a previously proposed model dubbed “epistatic capture” [102]. The authors reported that the combination of a MER20 and MER39 element form the enhancer and promoter of decidual prolactin in humans. By tracing the evolutionary history of these elements, they demonstrated that the MER39 element inserted nearby the preexisting MER20 element, after which 7 nucleotide substitutions added 4 TFBSs which together drove elevated expression of prolactin. The authors argue that the presence of nearby TFBSs caused the stabilization of motifs and maturation of proto-motifs residing within the MER39 element [102]. Our findings for RSINE1 are in agreement with this model and suggest that the cis-regulatory landscape surrounding the insertion site of a TE profoundly influences its mutational path and cis-regulatory trajectory.

An important implication of the model outlined above is that it may also promote the turnover of TFBSs and cis-regulatory elements during evolution. Our data for RSINE1 suggest that RSINE1 are more likely to acquire a CR BS when they inserted near an existing CR BS. This process, in turn, would lead to redundancy in CR-binding, which in some cases might lead to the mutational decay of the ancestral binding site, or be preserved by natural selection as a ‘buffer’ against harmful cis-regulatory variation [103]. Choudhary and colleagues recently reported an elegant demonstration of redundant TFBSs introduced by transposition [104]. The authors show that TE-derived CTCF sites inserted nearby existing CTCF sites in human and mouse reinforce and maintain conserved higher order chromatin structure by conferring redundancy to conserved CTCF sites. The motif gain we see in RSINE1 insertions nearby existing CR BS could represent binding site turnover events. It is difficult to rigorously assess this without information regarding BMAL1 binding in the liver of other species, but this is an idea worthy of further investigation.

## Conclusions

In summary, we propose that evolution of circadian regulatory elements from RSINE1 has followed this trajectory: (a) RSINE1 expanded in a single burst of rapid amplification in the murine lineage prior to the divergence of the mouse and the rat, (b) depending on the genomic context, most insertions were neutral or mildly deleterious due to their CR binding and mild regulatory activity in new genomic locations, and (c) a small subset of elements were tolerated or advantageous due to their proximity to existing circadian regulatory elements and have been co-opted as circadian enhancers either through binding site turnover, introduction of regulatory redundancy and buffering, or rarely by wiring a new gene into the circadian regulatory network (Fig. 6B).

Our study of gradual emergence of cis-regulatory activity within RSINE1 suggests an alternative to the copy-and-paste model of TE co-option whereby a TE inserts a ready-made enhancer module in a location where that activity becomes immediately beneficial [51,105–107]. While this is an attractive model, such cases are likely to be rare because TEs carrying such ready-made cis-regulatory modules, such as those found within the long terminal repeats of retrotransposons and endogenous retroviruses, are substantially more likely to have deleterious effects upon insertion than they are to be adaptive. Instead, TEs with little to no intrinsic cis-regulatory activity upon insertion are more likely to be tolerated near genes and existing cis-regulatory elements. The ‘low regulatory profile’ of these TEs might enable them to achieve higher copy number and disperse, on a large scale, proto-motifs poised for the emergence of new cis-regulatory elements. SINEs make excellent candidates for this model because they are short, noncoding, and typically derived from tRNA and other Pol III transcripts, thereby reducing their capacity to interfere with Pol II-regulation. Perhaps thanks to this property, SINEs have attained enormous copy numbers in virtually all mammalian genomes and they accumulate closer to genes than other types of TEs such as LINEs or ERVs [39,40,50,108]. It may not be coincidental that many studies have identified ancient SINEs co-opted as enhancers and other complex cis-regulatory elements [109–116], but it has been difficult to discern features of these elements that promoted their exaptation. We speculate that we have caught RSINE1 at an early stage of the co-option process and that the principles uncovered herein illuminate how seemingly benign TEs like SINEs have been a pervasive source of cis-regulatory elements in vertebrate evolution.

## Methods

### Reanalysis of sequencing data

Fastq files were downloaded from SRA (Table 1). Reads were trimmed and filtered with cutadapt [117] (arguments: -a AGATCGGAAGAGCACACGTCTGAACTCCAGTCAC; -A AGATCGGAAGAGCGTCGTGTAGGGAAAGAGTGTAGATCTCGGTGGTCGCCGTATCATT; --minimum-length=25), and aligned to the mm10 reference genome using bowtie2 [118] in paired end mode where applicable (arguments: --local; --very-sensitive-local; --no-unal; --no-mixed; --no-discordant; -I 10; -X 1000). SOLiD reads were aligned using bowtie [119] in colorspace (arguments: -C; -m 1). Alignment files were processed, sorted, and filtered for uniquely mapping reads (q > 1) using samtools [120] and PCR duplicate reads removed with Picard MarkDuplicates. Peaks were called using macs2 [121] (arguments: -g mm; --call-summits).

### Genomic interval analyses

All intersections of sets of genomic intervals were performed using bedtools intersect [122] (arguments: -wo; -f 0.5), which requires that 50% of a feature in the query file overlaps a feature in the subject file in order to record an intersection. Non-intersecting regions were separately determined (arguments: -v; -f 0.5). Sets of CR ChIP-seq peak coordinates were intersected as queries against the output of RepeatMasker [75], which was retrieved from the UCSC genome browser and filtered to remove low complexity regions and simple repeats. All calculations of distances between genomic intervals was calculated using bedtools closest (arguments: -d).

### Expression profiles and heatmaps

Normalized signal coverage files were generated using DeepTools bamCoverage [123] (arguments: --binSize 10; --normalizeTo1x 2150570000). Single end reads were extended to match the average library insert fragment size reported at publication; empirically determined fragment size was used when paired-end sequence data was available. The scaling normalization employed here presents coverage in each bin as a ratio of observed coverage to expected genome wide coverage assuming equal distribution of reads (1x coverage), and accounts for the mappable size of the genome, read depth, and read length. Profile meta-plots and heatmaps were generated using DeepTools [123]. Coverage matrixes were generated using computeMatrix in reference-point mode (arguments: --referencePoint center; --missingDataAsZero). Heatmaps were created with plotHeatmap and profiles were created with plotProfile (arguments: --plotType se). Quantification of signal in peaks was performed by using bigWigSummary to extract mean coverage at each genomic interval; followed by statistical testing in R using the packages kruskal.test and dunn.test. For visualization purposes, when plotting heatmaps designed to compare between sets of intervals with different sample sizes (for example, RABS-RSINE1 n=328 and Non-RABS-RSINE1 n=115,806), the larger set of intervals was subsampled to the match the smallest set of intervals prior to matrix computation.

### Gene Ontology Analysis

Enrichment of mouse phenotypes with each group of genomic intervals was calculated by GREAT [77] using the two nearest genes within 1 Mb of each E-Box motif.

### Repeat element enrichment

Enrichment was calculated as described [78] (https://github.com/4ureliek/TEanalysis) by shuffling TEs such that the distance to the nearest TSS and number of elements on each chromosome was maintained. These shuffled TEs were intersected with the mm10 RepeatMasker annotations [75] to determine the number of expected intersections from each TE and TE family. This procedure was bootstrapped 1000 times. A binomial test was then performed to determine significance, comparing the average expectation to the observed value.

### Consensus Alignment

Consensus sequences of RSINE1, B4, and B4A were obtained from Repbase [74] and aligned using Clustal Omega [124]. The resulting alignment was visualized and manually curated using Jalview [125]. Relevant motifs were identified manually.

### Motif analysis

Sequences of bound, OC unbound, and all RSINE1 elements were retrieved using bedtools getfasta [122]. Unbiased motif discovery was performed using DREME [126] with bound RSINE1s as primary sequences, and both OC unbound and all RSINE1s as control sequences, sequentially. This procedure was repeated using the 500 bp flanks (left and right) of each RSINE1 element in each set. Locations of E-box (CACRTG), RORE (RGGTCA), and D-Box (TTATGYAA) motifs were predicted genome wide using FIMO [127] (arguments: --thresh 0.01; --max-stored-scores 10000000). Bedtools merge [122] was used to merge overlapping motifs and average the FIMO score over the merged region, and the utility bedGraphToBigWig was used to generate coverage tracks where the coverage represents genome wide motif occurrence weighted by score. Heatmaps were generated as above. The distance to the nearest E-Box motif from each set of intervals (peaks and repeats) was calculated using bedtools closest [122] (arguments: -d), and E-Boxes which fall within the boundaries of the peak (and therefore are distance = 0) were retained. For analysis of distance between motifs, we filtered coordinates of predicted motifs by FIMO log-odds score, retaining only motifs with the highest possible score. Bedtools intersect (arguments: -f 1; -wa) reported motifs that completely overlap with a given set of repeats; the distance from these overlapping motifs to the nearest non-overlapping motif was calculated using bedtools closest (arguments: -io; -d).

### Conservation analysis

We attempted to map sets of genomic regions from the mm10 genome to the hg38 or rn6 genomes using liftOver (Kuhn et al., 2013) using appropriate chains, and counted the fraction of intervals which did or did not lift over from each set of elements.

### RSINE1 phylogeny

We began with sequences of the 328 bound RSINE1 elements, as well as 1000 randomly selected elements from the set all RSINE1s in the mm10 genome assembly (n=115,806). We aligned these sequences using Clustal Omega [124] with default settings. From this alignment, we calculated a maximum likelihood tree with FastTree 2 [128] using default settings for nucleotide alignments. The tree was visualized with ggtree [129] and branches were colored by their designation as bound or unbound RSINE1 elements.

### Mouse/Rat Motif Comparison

We started with the set of CR bound RSINE1 elements (n=328) and unbound elements residing in open chromatin (n=4583) and attempted to assign syntenic orthologs of each element using liftOver [84]. We were able to find orthologous regions for 195 bound RSINE1 elements (59.5%) and 2662 unbound elements (58.1%). We queried the sequences of each of these elements using bedtools getfasta [122] from the mm10 and rn6 genome assemblies, and used DREME [126] as above to find motifs enriched in orthologous bound RSINE1 elements when compared to unbound RSINE1 elements.

### Tissue Specific BMAL1 Analysis

We started with coordinates of BMAL1 binding sites in the liver, kidney, and heart [72] in the mm10 genome assembly. We performed an intersection as described above (requiring 50% overlap of a given peak with a repetitive element to designate that peak as a RABS) between each of these sets of coordinates and the output of RepeatMasker [75] which was retrieved from the UCSC genome browser and filtered to remove low complexity regions and simple repeats. We counted the number of Non-RABS, RABS, and RSINE1-RABS and plotted the fraction each category represents with respect to the total set of peaks, and the fraction of RABS which are RSINE1-derived.

### Luciferase Reporter Assays

We accessed the RSINE1 consensus sequence from Repbase [74] and created an “*in silico* evolved” version by changing a total of 7 nt to “perfect” the E-Box and RORE motifs highlighted in Fig. 4A (RORE 1: unchanged; RORE 2: AGTTCG>AGGTCA; RORE 3: AGGAGA>AGGTCA; E-Box 1: CACATG>CACGTG; E-Box 2&3: CACGCG>CACGTG). We also accessed, from the mm10 genome assembly, the sequences of *Tmem267* and *Aaed1* RSINE1 elements highlighted in Fig. S6, both with and without 100 nt of flanking sequence on the 5’ and 3’ ends. All of these sequences were synthesized as gBlocks® Gene Fragments (Integrated DNA Technologies, Coralville, IA, USA) and cloned into pGL3::PRO (Promega Corporation, Madison, WI, USA) downstream of the luciferase gene. To generate vectors of *Tmem267* and Aaed1 RSINE1 flanking sequence only, existing vectors were amplified by PCR using Q5 High-Fidelity DNA Polymerase (New England Biolabs, Ipswich, MA, USA) with primers designed to delete the RSINE1 element by site directed mutagenesis. All vectors were validated by Sanger sequencing. Hepa 1-6 cells were obtained from ATCC and were subject to co-transfection with 500 ng of each of the constructs described above in conjunction with 500 ng pRL::SV40 Renilla Luciferase Control Vector (Promega Corporation, Madison, WI, USA) using Lipofectamine-2000 (Thermo Fisher Scientific, Waltham, MA, USA). After 18h, firefly and Renilla luciferase luminescence was measured using the Dual-Glo® Luciferase Assay System (Promega Corporation, Madison, WI, USA). Firefly luminescence values were normalized to Renilla luminescence and compared to the empty pGL3::PRO vector.

## Declarations

### Ethics approval and consent to participate

Not Applicable.

### Consent for publication

Not Applicable.

## Availability of data and materials

All data used in this study was previously published and accession numbers can be found in Table 1.

## Authors’ contributions

CF, JJ, and HS conceptualized the study. JJ and HS performed genomic data analysis. JJ conducted reporter assays. JJ, CF, and HS wrote the manuscript. All authors read and approved the final manuscript.

## Acknowledgements

The authors acknowledge Tianyi Zhang and Fanliu Kong for technical assistance with cloning, and thank members of the Feschotte Lab and Lis Lab for helpful discussions.

## Supplemental Figures

**Figure S1:**
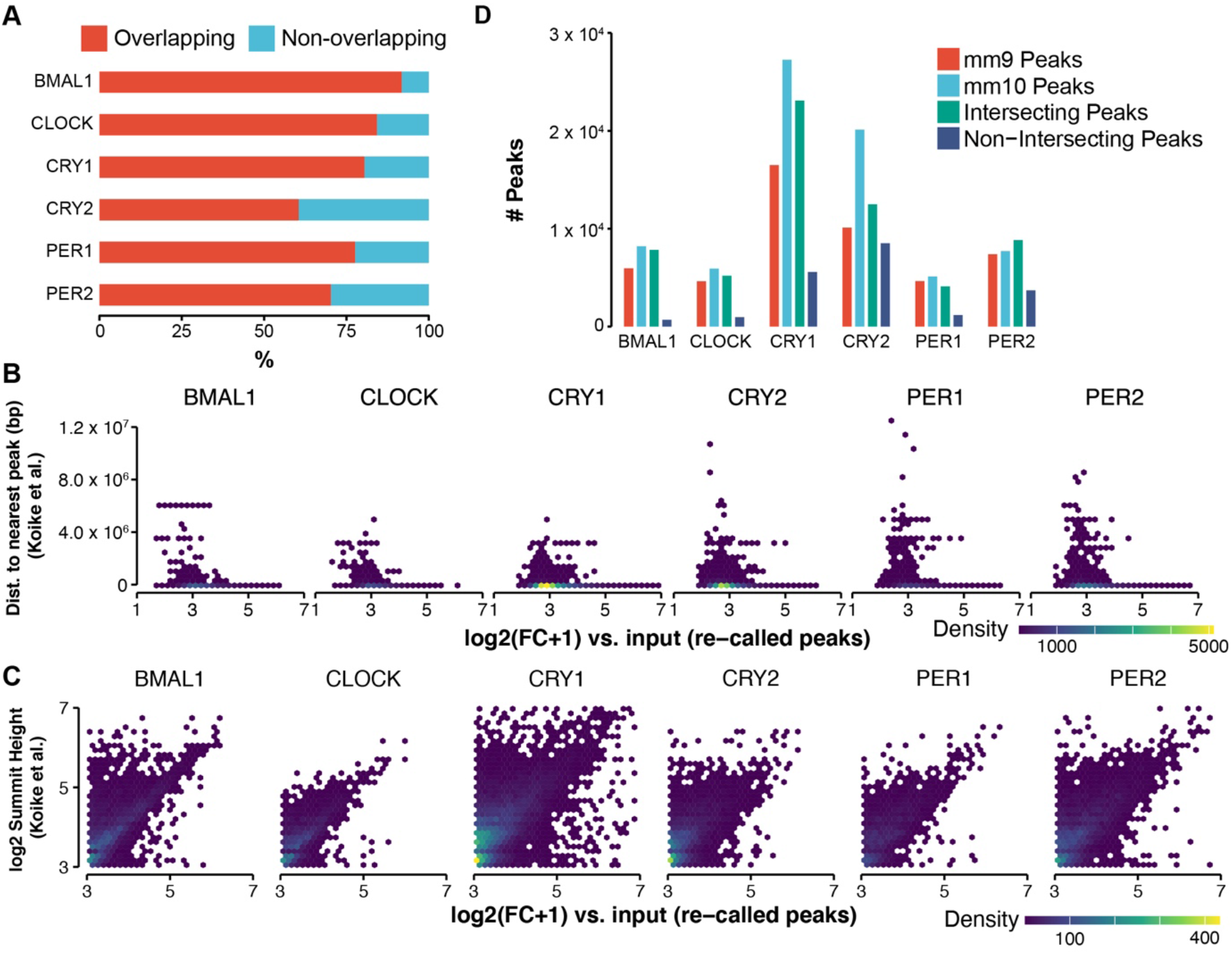
New analysis of Circadian Regulator ChIP-seq data is similar to published results. **(A)** Percentage of peaks determined in the new analysis described in this report that overlap with peaks in the published set. **(B)** Fold-change as reported by macs2 in the re-analysis compared to summit height for each peak as reported in the published set of peaks. **(C)** Fold-change as above for each peak in the new analysis compared to the distance to the nearest peak in the published set (peaks in the published set were mapped to mm10 coordinates using LiftOver). **(D)** Number of peaks in the published set and in the new analysis presented here for each CR, as well as the number of peaks which are shared or not shared between the two datasets. For **(C–D)**, the plot space was divided into hexbins and color-scaled according to the density of points in each bin to accommodate over-plotting.

**Figure S2:**
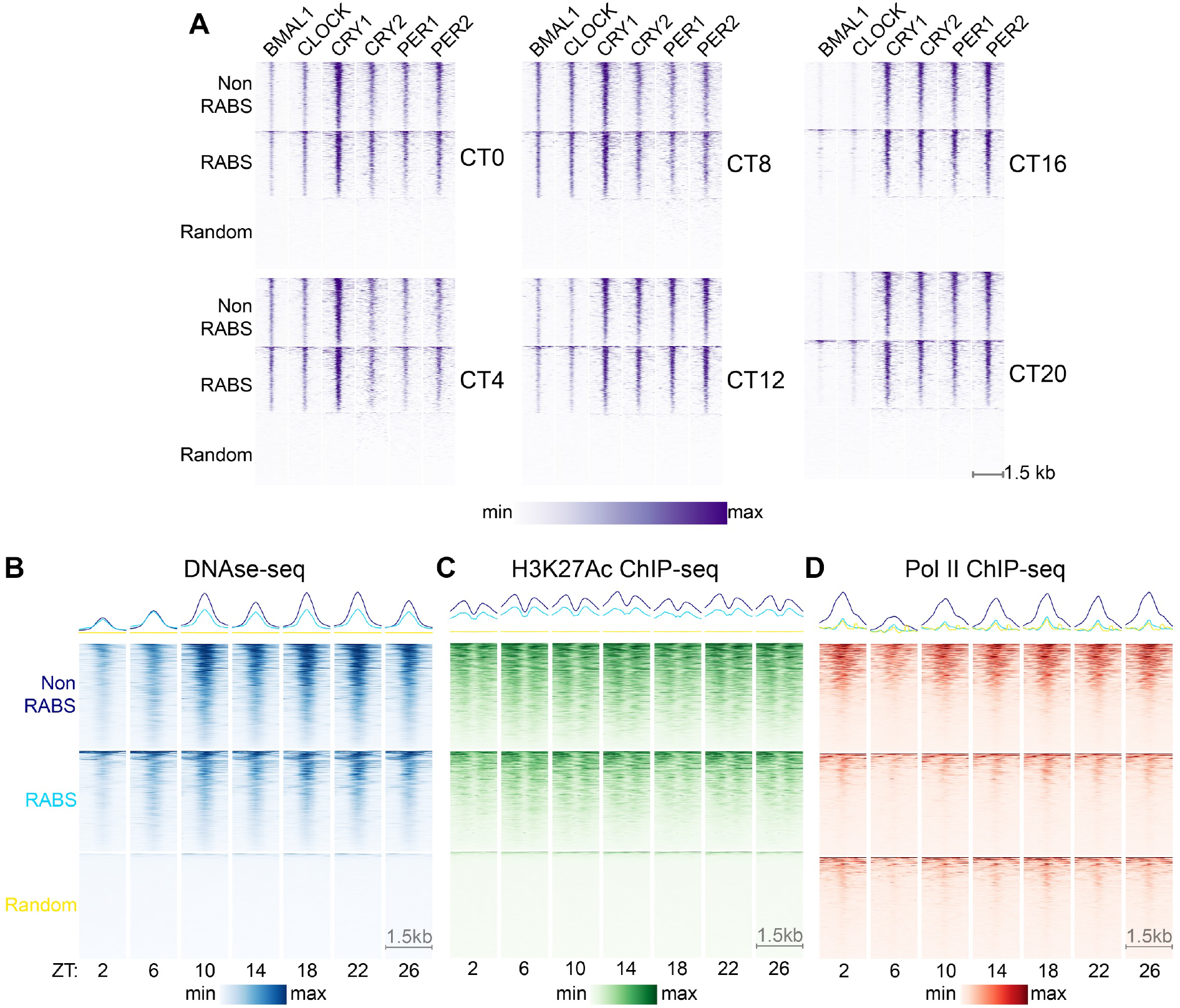
RABS have a similar pattern of CR occupancy but different chromatin properties compared to Non-RABS. **(A)** ChIP-seq signal of each CR centered at BMAL1 ChIP-seq peaks at RABS (n=1014), randomly selected Non-RABS (n=1014), and randomly selected repeats matching the familial composition of RABS (n=1014) over Circadian Time (CT). **(B)** DNase-seq signal over Zeitgeber time (ZT). Regions as in **(A)**. **(C)** H3K27Ac ChIP-seq over ZT. Regions as in **(A)**. **(D)** Pol II ChIP-seq at over ZT. Regions as in **(A)**. In all panels, heatmaps are sorted by mean signal intensity across all rows in a block of eatmaps. Color scaling is min-max within each block.

**Figure S3:**
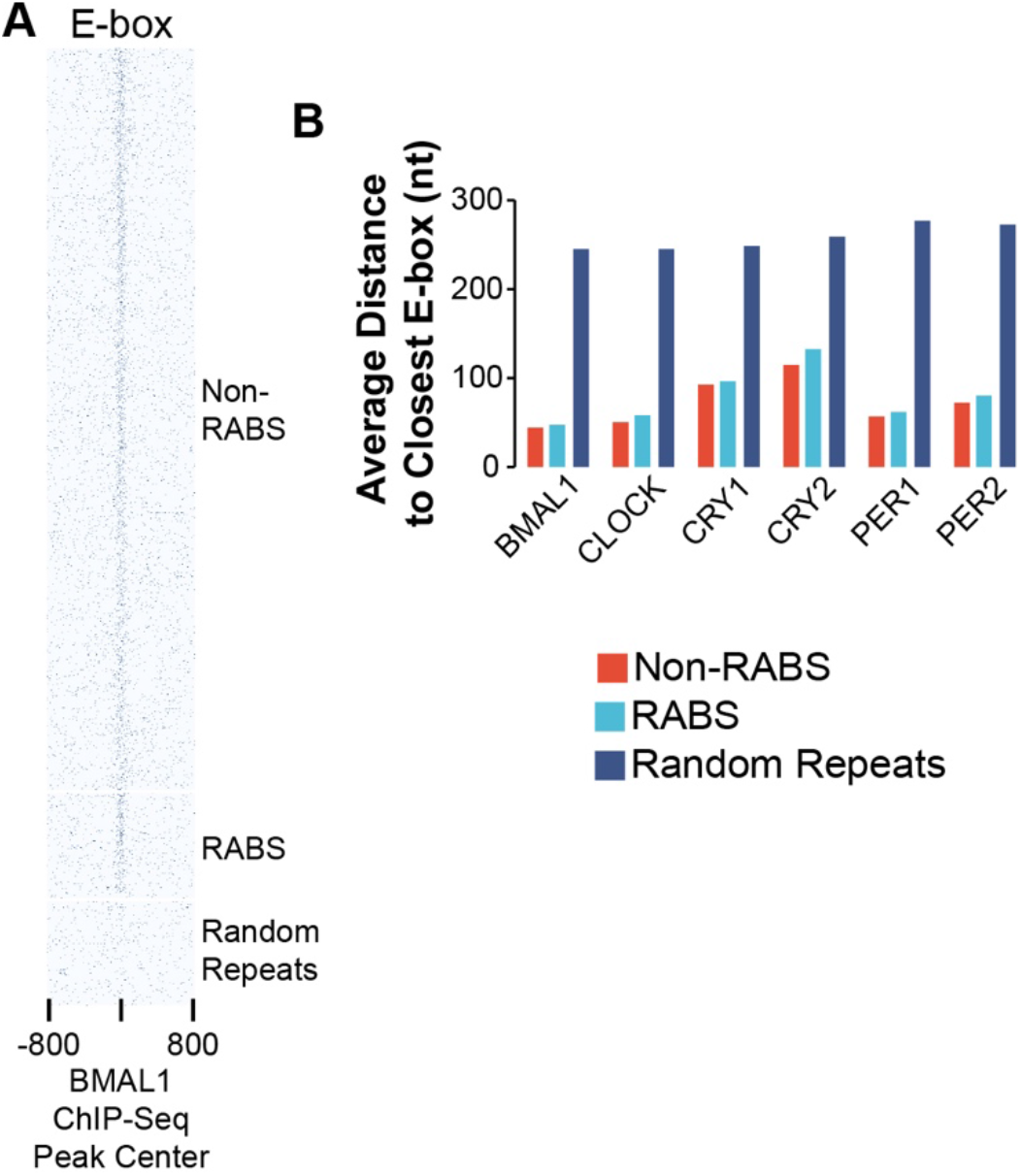
Repeats contribute E-Box motifs. **(A)** Occurrence of E-Box motifs at RABS and Non-RABS BMAL1 ChIP-seq peaks, as well as a set of repeats randomly selected to match the familial composition of repeats associated with BMAL1 RABS. Signal strength (color intensity) is relative to motif similarity to the consensus motif. **(B)** Average distance to the closest E-Box motif from RABS or Non-RABS ChIP-seq peak centers as well as a randomly selected set of repeats that matches the familial composition of each set of CR RABS.

**Figure S4:**
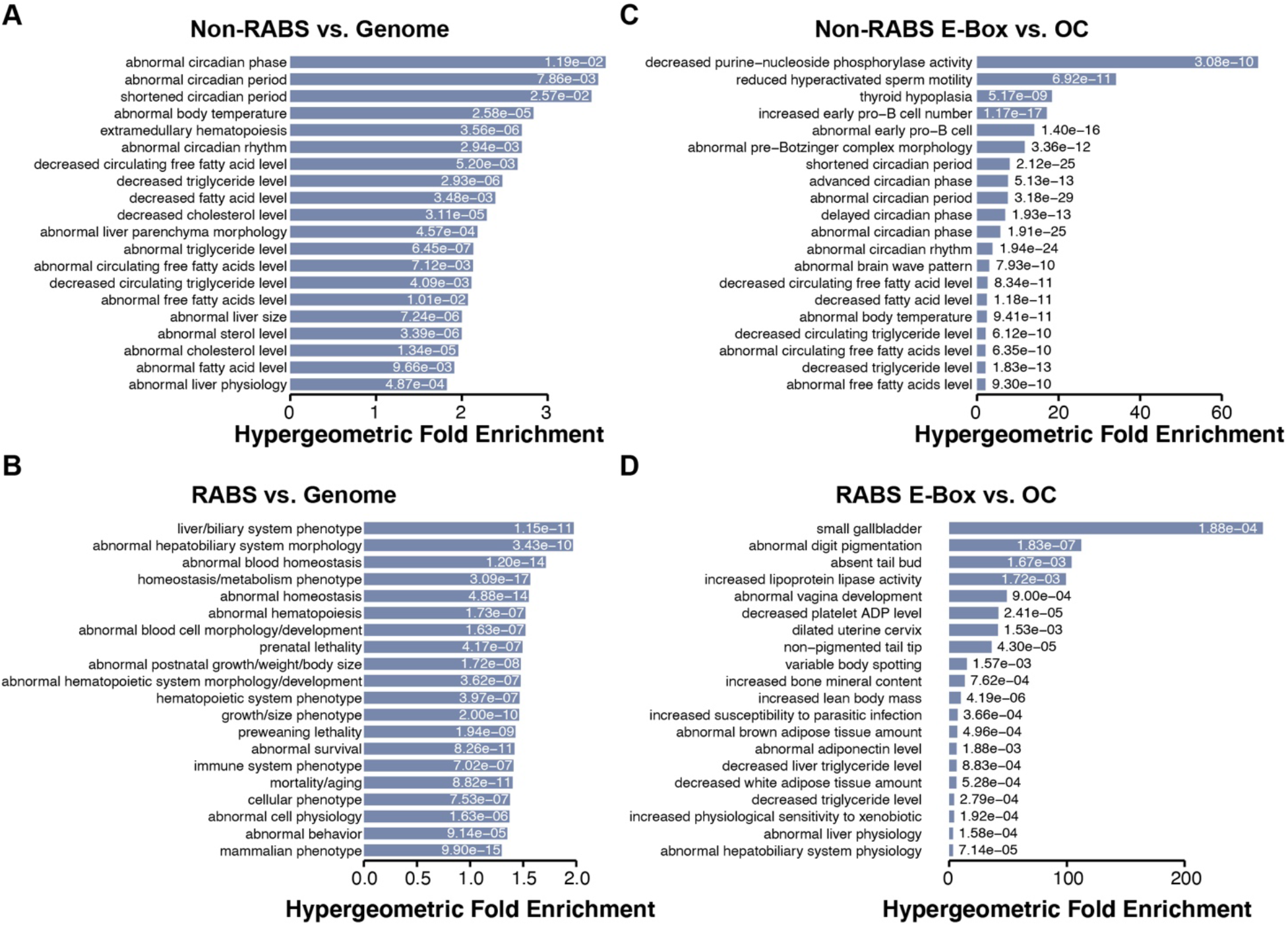
RABS-proximal genes are enriched for more lineage-specific phenotypic associations than Non-RABS-proximal genes. **(A)** Phenotypic association enrichment of Non-RABS E-Box motifs using the entire mouse genome as background. **(B)** Phenotypic association enrichment of RABS E-Box motifs using the entire mouse genome as background. **(C)** Phenotypic association enrichment of Non-RABS E-Box motifs using open chromatin regions defined by DNase accessibility at any timepoint as background **(D)** Phenotypic association enrichment of RABS E-Box motifs using open chromatin regions defined by DNase accessibility at any timepoint as background Erichments were calculated by the tool GREAT using the two nearest genes within 1 Mb of each x motif. FDR corrected hypergeometric q-values are either overlaid or adjacent to each bar.

**Figure S5:**
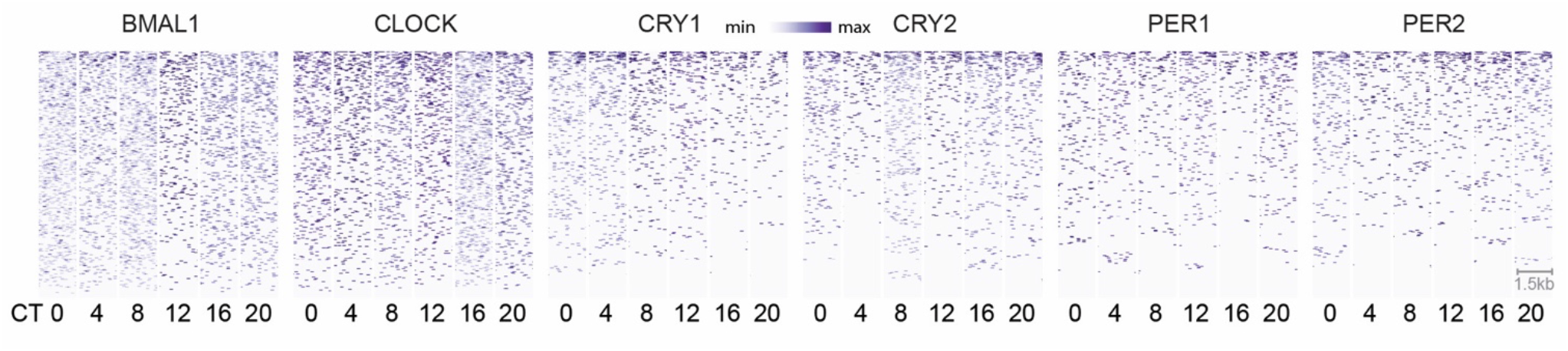
RSINE1 elements not in open chromatin do not display oscillatory CR ChIP-seq signal. ChIP-seq signal for each CR across circadian time (CT) at a randomly selected set of 328 RSINE1 elements which are not bound by CRs and do not reside in open chromatin regions as defined by DNase-seq. Heatmaps are centered at the middle of each element and extend 750 bp in each direction, and are sorted by mean signal intensity across all rows for a given CR. Color scaling is relative to min-max signal within each block of heatmaps.

**Figure S6:**
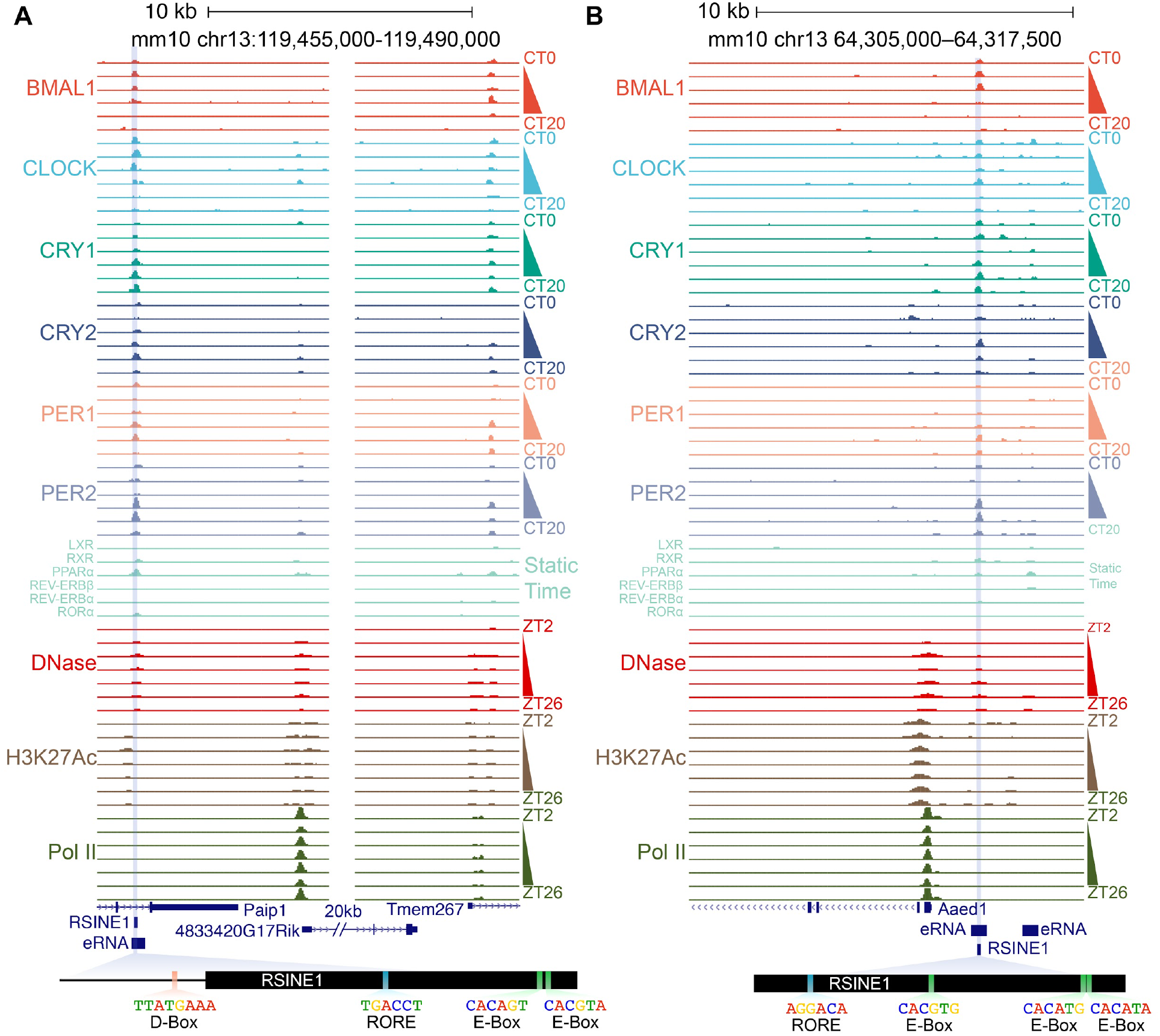
UCSC genome browser screenshots of the two RSINE1s selected for luciferase reporter assays. **(A)** Browser shot of the *Tmem267* locus. **(B)** Browser shot of the *Aaed1* locus. In each panel, coordinates of oscillating eRNAs are shown alongside the location of an overlapping CR bound RSINE1. The location of the RSINE1 cloned into the luciferase reporter in Fig 7 is highlighted, and the motif content of that particular element is shown. The *Tmem267* RSINE1 shows a small upstream region where a D-Box motif was found adjacent to the insertion site.

